# On the Effect of Intralocus Recombination on Triplet-Based Species Tree Estimation

**DOI:** 10.1101/2021.11.06.467557

**Authors:** Max Hill, Sebastien Roch

**Affiliations:** Department of Mathematics, University of Wisconsin–Madison

## Abstract

We consider species tree estimation from multiple loci subject to intralocus recombination. We focus on *R*\*, a summary coalescent-based methods using rooted triplets. We demonstrate analytically that intralocus recombination gives rise to an inconsistency zone, in which correct inference is not assured even in the limit of infinite amount of data. In addition, we validate and characterize this inconsistency zone through a simulation study that suggests that differential rates of recombination between closely related taxa can amplify the effect of incomplete lineage sorting and contribute to inconsistency.

## 1 Introduction

Species tree estimation from genomic data is complicated by various biological phenomena, among them hybridization, horizontal gene transfer, gene duplication and loss, and incomplete lineage sorting (ILS) [SS20]. ILS in particular is a source of phylogenetic conflict, in which gene trees and species tree do not share the same topology, especially for species trees with short internal branches [SS20]. Of some interest is the existence of an anomaly zone for species trees, in which the most probable topology in the gene tree distribution differs from the topology of the species tree [DR06, DR09, Deg13] (see also [RELY20, BH20] for a more recent discussion of these and other relevant issues).

The existence of an anomaly zone has served as an impetus for the development of summary coalescent-based methods, such as *R*\*, MP-EST, BUCKy, ASTRAL, and others [DDBR09, LYE10, LKDA10, MRB^+^14]. Some of these methods are based on the facts that rooted triples and unrooted quartets are special cases in which no anomaly zone exists [Deg13, KYT20] and also provide sufficient information to reconstruct the full phylogeny [SS^+^03, Ste16]. Provided that the gene trees are estimated without error, such methods can provide statistically consistent methods of estimating species tree topology [War17].

A common assumption of coalescent-based models based on the multispecies coalescent (MSC) [RY03, RELY20] is that recombination occurs between genes (or loci)—so that gene trees may be assumed unlinked or statistically independent—but that *intralocus recombination* (i.e., recombination occurring *within* gene sequences), does not occur [EXJ^+^16, BH20]. The significance of the latter assumption—that is, the impact of intralocus recombination on phylogenetic inference—is a matter of present interest [ZY21, BH20] and much debate about its significance when unaccounted for [LK12, SG18, EXJ^+^16]. One justification for assuming no intralocus recombination is that within-gene recombination may break gene function [SS20].

An influential simulation study argued that even high levels of intralocus recombination do not present a significant challenge for species tree estimation relative to other biological phenomena [LK12]. On the other hand, the authors of [SG16] suggest the absence of intralocus recombination may be an unreasonable assumption in real data, such as protein-coding genes in eukaryotes [MLH19, BH20], and particularly in the case of species phylogenies with many taxa [SG18]. In particular, the potential for intralocus recombination to distort gene tree frequencies has been recognized as a challenge to summary coalescent-based methods, and [LK12] has been critiqued for its focus on shallow divergences and limitation to a low number of loci and taxa [SG18].

In this paper we take an analytical approach to investigate the effect of intralocus recombination. We prove that intralocus recombination has the potential to confound *R*\*, a summary coalescent-based methods based on inferring rooted triples. That is, we show that correct inference of rooted triplets cannot be guaranteed in the presence of intralocus recombination, assuming a distance-based approach is used for gene tree reconstruction. We then present a simulation study which characterizes the “inconsistency zone”, i.e. the regime of parameters for *S* in which rooted triple inference does not converge to *S* as *m* → ∞. We find that the effect arises when differential rates of recombination are exhibited between closely-related taxa. We also discuss related implications for quartet-based methods.

### 1.1 Key Definitions

A *species phylogeny* 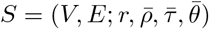 is a directed binary tree with vertex set *V*, edge set *E*, root *r* ∈ *V*, and *n* labeled leaves *L* = [*n*], such that each edge *e* ∈ *E* is associated with a length *τ*_*e*_ ∈ (0, ∞), expressed in coalescent units, a recombination rate *ρ*_*e*_ ∈ [0, ∞), and a mutation rate *θ*_*e*_ ∈ [0, ∞). It is assumed that there exists an ancestral population common to all leaves of *S*, i.e., a population above the root, with respective mutation and recombination parameters. In this paper, mutation rates are assumed to be per site per coalescent unit (a coalescent unit being 2*N*_*e*_ generations for diploid organisms, where *N*_*e*_ is the effective population size); the recombination rates are per locus per coalescent unit.

The general question considered here is how to reconstruct the topology of the species phylogeny from gene sequence data sampled from its leaves. This sequence data takes the form of multiple sequence alignments; a *multiple sequence alignment* (MSA) is an *n* × *k* matrix *M* whose entries are letters in the nucleotide alphabet {*A, T, C, G*} such that entries in the same column are assumed to share a common ancestor. The phylogenetic reconstruction problem in this paper is to recover the topology of *S* from *m* independent samples of *M*.

A *rooted triple* is a rooted binary phylogenetic tree with label set of size three; we use the notation *XY* |*Z* (or equivalently *Y X* |*Z*) to denote a rooted triple with leaves *X, Y, Z* having the property that the path from *X* to *Y* does not intersect the path from *Z* to the root [SS^+^03]. The term *species triplet* refers to a restriction of *S* to three of its leaves. A rooted triple *XY* |*Z* is said to be *uniquely favored* if it appears in more gene samples than either of the other two rooted triples *XZ*|*Y* or *Y Z*|*X*.

### 1.2 Inference Methods

This paper considers *Majority-Rule Rooted Triple*, or *R*\*, a consensus-based pipeline for species tree estimation. *R*\* utilizes the fact that the full topology of *S* is uniquely determined by, and hence can be recovered from, its rooted triples [Ste16]. The *R*\* pipeline has three steps: first, for each gene, infer a rooted triple for each triplet of leaves *X, Y, Z* ∈ *L*. Second, make a list of uniquely favored triples from the *m* sampled genes. Finally, construct the most-resolved topology containing only uniquely favored triples. When gene trees are drawn independently according to the MSC, it holds that for every set of three taxa, the most probable rooted triple in the gene tree distribution matches the rooted triple obtained by restricting the species tree *S* to that set of three taxa; for this reason, the topology of the *R*\* consensus tree converges to that of *S* [DDBR09].

Since we are interested in the inference of the species-tree topology from *sequence data*, we consider a distance-based approach in which a species triplet with leaves *X, Y, Z* is inferred to have topology *XY* |*Z* if

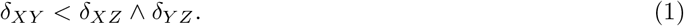

where *δ*_*XY*_ = *δ*_*XY*_ (*M*_*k*_) is the number of mismatching nucleotides between sequences **s**_*X*_ and **s**_*Y*_ (*X, Y* ∈ *L*). We refer to this inference procedure as ***R***\* **with sequence distances**.

### 1.3 Multispecies Coalescent with Recombination

The model considered here, which we term the *Multispecies Coalescent with Intralocus Recombination*, or MSCR, uses the ancestral recombination graph (ARG) model from [GM97] (see also [Are13]) within the framework of the multi-species coalescent (MSC) [RY03, RELY20, DR09]. In the single-population ARG [GM97], ancestors are represented by edges in the graph (see Figure 1a), and the number *N* of ancestors, or *gene lineages*, at time *t* is a bottom-up birth-death process in which births (recombination events) occur at rate *ρN* and deaths (coalescent events) occur at rate *N* (*N* − 1)/2. When a coalescent event happens, two edges are chosen at random and merged into one. When recombination occurs, a randomly chosen lineage splits into two parent lineages. Each recombination vertex is labeled by a number *b*, chosen uniformly on [0,1]; this number is the *breakpoint* of the recombination.

**Figure 1:**
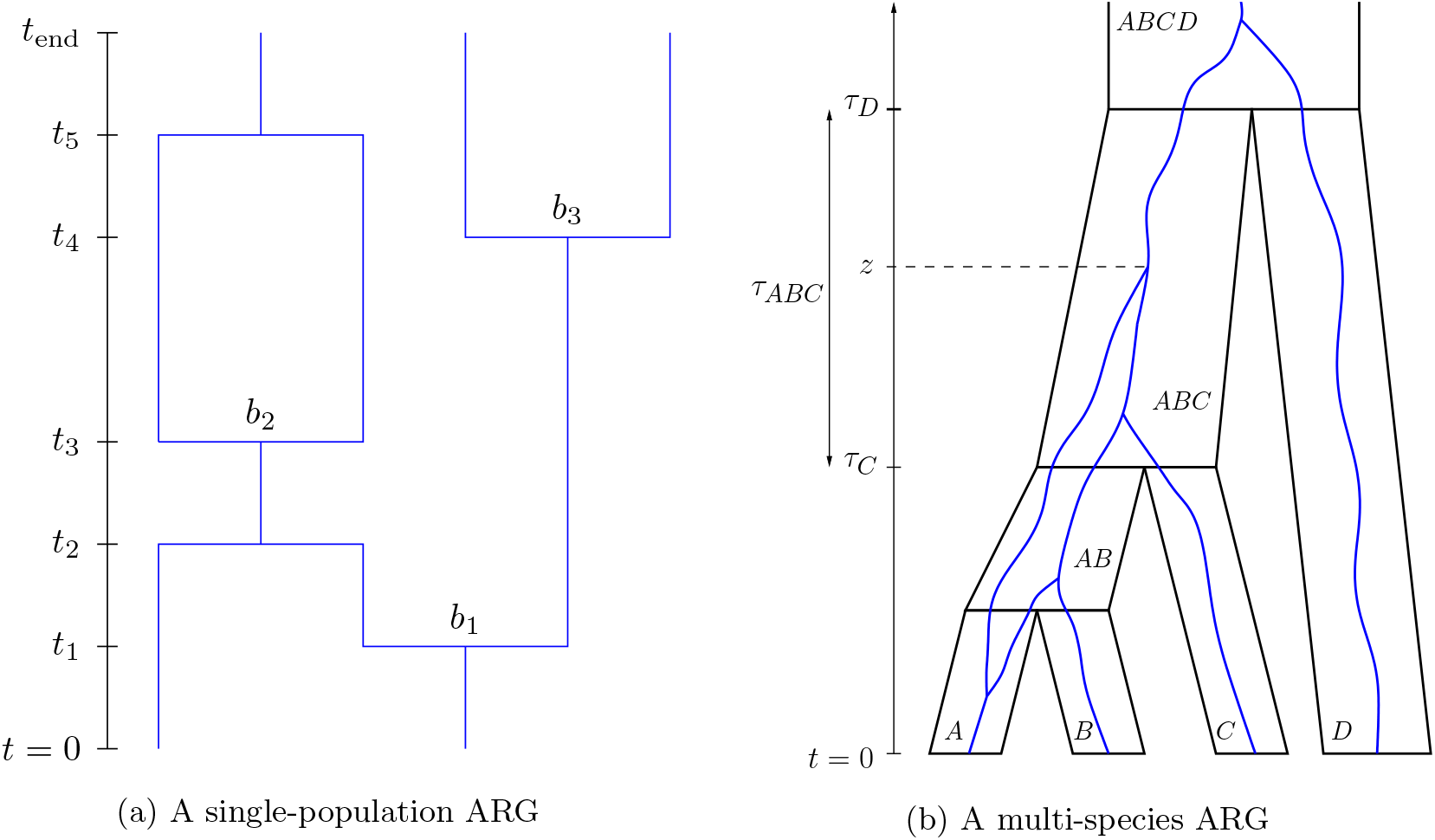
Two depictions of an ARG, in a single population (left) and in the multispecies case (right). In Figure 1a, two lineages enter the population at time 0 and three exit at time *t*_end_. Coalescent events occurred at times *t*_2_ and *t*_5_. Recombinations with breakpoints *b*_1_, *b*_2_, *b*_2_ occurred at times *t*_1_, *t*_3_, and *t*_4_. Figure 1b depicts a multispecies ARG (in blue) within the four-taxa tree *S* considered in Corollary 1.

The single-population ARG can be extended to multiple species in a manner similar to the MSC: at time *t* = 0, each leaf of *S* begins with a single lineage, and these lineages evolve in a bottom-up manner according to the ARG process along each edge of a fixed species tree (see Figure 1b). If 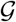 is a rooted directed graph with edge lengths and leaf and breakpoint labels obtained in this manner, then we say that 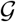 is **generated according to the MSCR process on *S***. In this scheme, the locus is modeled by the unit interval, and for each site *x* ∈ [0, 1], a *marginal gene tree* 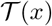 can be obtained by tracing upward along the edges of 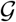 starting from the leaves; if a recombination vertex is reached with breakpoint *b*, take the left path if *x* ≤ *b* and the right path if *x* > *b*. This yields a rooted edge-weighted binary trees; a simple example is shown in Figure 2. The set of marginal gene trees 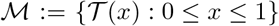 is almost surely finite [GM97]. For each 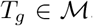, define 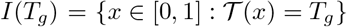, and define *w*_*g*_ = |*I*(*T*_*g*_)|, where |·| denotes Lebesgue measure. In words, *I*(*T*_*g*_) is the identical-by-descent segment of the locus having genealogy *T*_*g*_, and *w*_*g*_ is the proportion of sites with genealogy *T*_*g*_.

**Figure 2:**
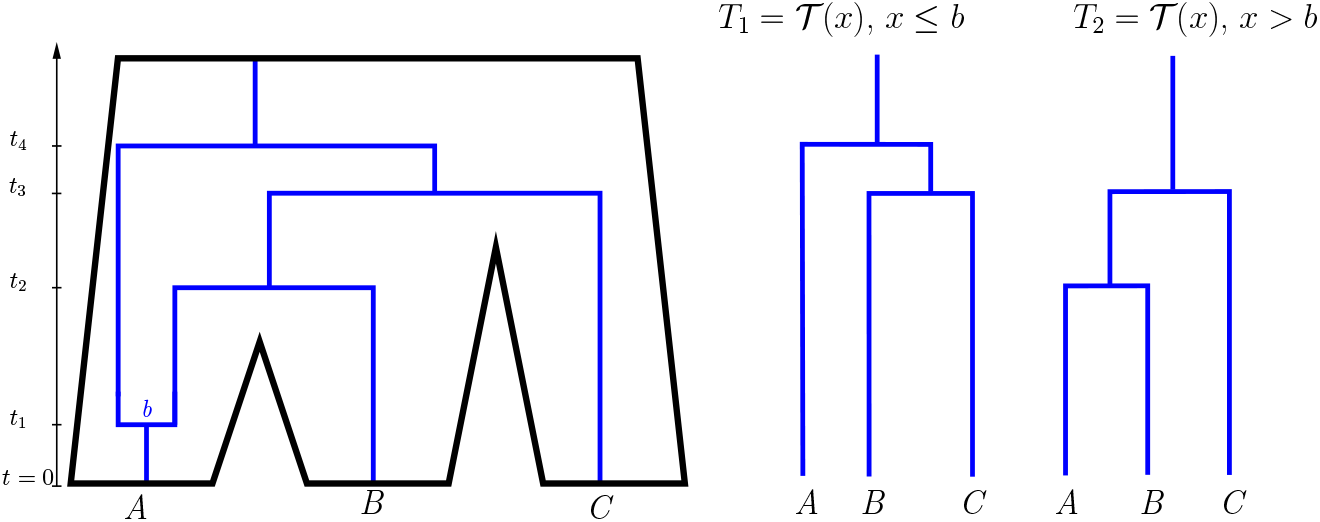
On the left, an ancestral recombination graph (in blue) is shown within a 3-taxa tree *S* (in black). The times of coalescence and recombination events are labeled *t*_1_, … *t*_4_ on the time axis, and the breakpoint associated with the recombination event is labeled *b* ∈ [0, 1]. On the right, the corresponding marginal gene trees *T*_1_ and *T*_2_ are shown.

Measuring time in coalescent units, this paper assumes that the per-site mutation rate is given by a fixed number *θ* > 0 which does not vary on *S*. For each *x* ∈ [0, 1], site *x* evolves independently according to the Jukes-Cantor process [JC^+^69, Ste16] on the tree 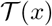. A somewhat more general description of this algorithm can be found in [DMNR17].

Thus, to model the evolution of a genetic locus consisting of *k* sites in which recombination breakpoints are distributed uniformly between them, a two-step process is followed. First, a multispecies ARG 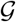 is generated according to the MSCR process on *S*, from which a marginal gene tree 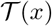 is obtained for each *x* ∈ [0, 1]. Second, for each *x* ∈ [0, 1] the Jukes-Cantor process is run with input tree 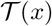 in order to generate a nucleotide 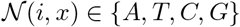 for each *i* ∈ *L*. The MSA *M*_*k*_ is then defined as the *n*×*k* random matrix with rows **s**_1_, … , **s**_*n*_ where for each *X* ∈ [*n*], **s**_*X*_ = (*s*_*X*_ (1), … , *s*_*X*_ (*k*)) where 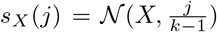, *j* = 0, 1, … , *k* − 1. In this case, we say that *M*_*k*_ is **generated according to the MSCR-JC(k) process on *S***. The phylogenetic reconstruction problem considered here can then be stated precisely as follows:

**Problem:** Let *S* be a species phylogeny with leaf labels *L* = [*n*]. Fix *k* ≥ 2. Given *m* independent samples 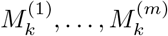, each generated according to the MSCR-JC(k) process on *S*, recover the topology of *S*.

### 1.4 Estimating Sequence Distances

Let 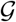 be generated according to the MSCR process on *S*, and 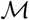 the corresponding set of marginal gene trees. Given a marginal gene tree 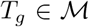, let 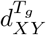 be the *evolutionary distance* between leaves *X* and *Y* on *T*_*g*_, defined as the expected number of mutations per site along the unique path between *X* and *Y*. It follows from the assumptions about the mutation process that 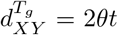, where *t* is the time of the most recent common ancestor of *X* and *Y* on *T*_*g*_. For example in Figure 2, 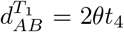 and 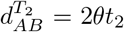. Define the *breakpoint-weighted uncorrected distance* by

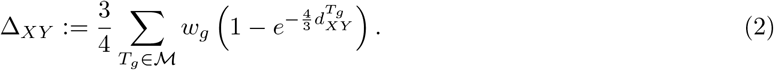

This formula, due to [Wan17], generalizes the uncorrected Jukes-Cantor distance to the setting of intralocus recombination; if no intralocus recombination occurs, then the right-hand side has only a single summand and reduces to the inverse of the Jukes-Cantor distance correction formula for a single non-recombining locus.

Our first lemma shows that *δ*_*XY*_ can be approximated by *k*Δ_*XY*_ when *k* is large.

#### Lemma 1.

*If M*_*k*_ *is generated according to the MSCR-JC(k) process on S then for all X, Y* ∈ *L*(*S*), *conditioned on* 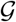, *δ*_*XY*_ (*M*_*k*_) = *k*Δ_*XY*_ + *o*(*k*) *almost surely as k* → ∞.

See Appendix A for a proof.

## 2 Inconsistency of R*

### 2.1 Statement and Overview

The main result is the following:

#### Theorem 1.

*For k sufficiently large, R*\* *using sequence distances is not statistically consistent under the MSCR-JC(k) model. That is, there exists a species phylogeny S such that the topology of the output of R*\* *using sequence distances does not converge in probability to the topology of the species tree.*

We discuss implications for quartet-based methods in Appendix C.

To prove Theorem 1, it suffices to consider a species tree *S* with *L* = {*A, B, C*} and topology *AB|C*. Denote edges of *S*, or *populations*, by the letters *A, B, C, AB,* and *ABC* as depicted in Figure 3 where *A, B, C* correspond to the leaf populations, *AB* is the parent edge of *A* and *B*, and *ABC* is edge extending above the root. The key idea is to allow recombination only in population *A*. In order to keep the analysis tractable, the recombination rate and length of edge *A* are chosen so that with high probability the number of recombinations is 0 or 1, so that the number of lineages on the ARG exiting population *A* (backwards-in-time) is either one or two. By choosing the internal branch length *τ*_*AB*_ sufficiently small, ILS occurs along that edge with high probability, so that all coalescent events on the ancestral recombination graph occur in the root population *ABC*. In that case, as long as the mutation rate is not too large, we show that, on the event *R*_1_*C*_0_ (see Figure 3), taxa *B* and *C* are more likely to be inferred as more closely related than taxa *A* and *B*, so that *R*\* converges to the wrong topology *BC|A* as the number *m* of samples grows.

**Figure 3:**
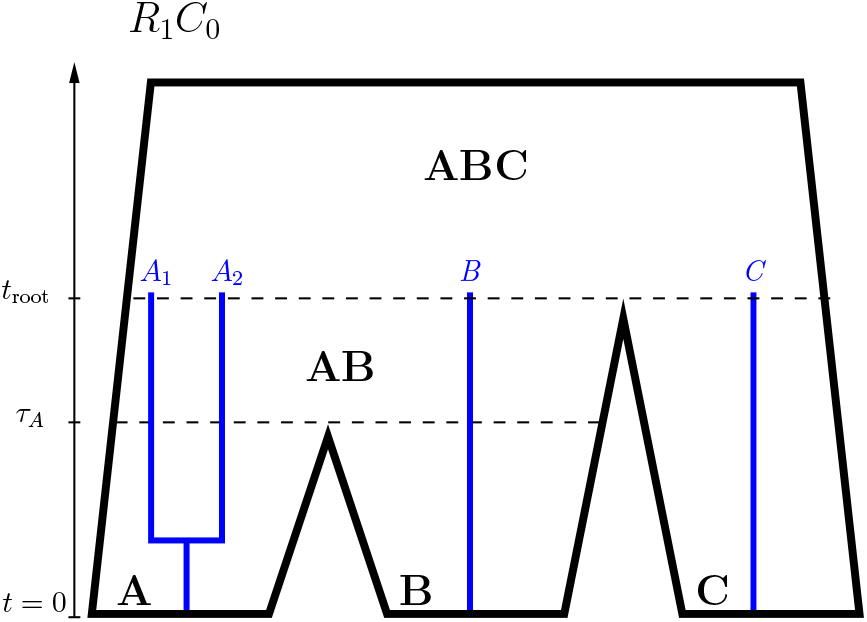
A depiction of the event *R*_1_*C*_0_. The portion of the ancestral recombination graph more ancient than *t*_root_ is not shown.

The mutation rate *θ* is assumed to be the same in all populations. The vector of recombination rates 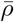 is defined by setting *ρ*_*A*_ = *ρ* > 0 and *ρ*_*X*_ = 0 for all *X* ≠ *A*. Assume *S* to be ultrametric. The populations *A* and *B* have length *τ*_*A*_ = *τ*_*B*_ > 0, the internal population *AB* has length *τ*_*AB*_, the age of the root *t*_root_ is given by *t*_root_ = *τ*_*A*_ + *τ*_*AB*_ = *τ*_*C*_. For now assume that *τ*_*AB*_ > 0 and *τ*_*A*_ > 0; their precise values will be determined later in the proof.

Let *M*_*k*_ be generated according to the MSCR-JC(k) process on *S*, and let *E*_*XY*|*Z*_ be the event that the rooted triple inferred from *M*_*k*_ using (1) is *XY*|*Z*. The following lemma implies that to prove Theorem 1, it will suffice to prove

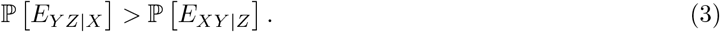

The *consistency zone* for *R*\* with sequence distances under the MSCR-JC(k) model is the set of species phylogenies *S* such that the topology of the *R*\* consensus tree converges in probability to the topology of *S* as *m* → ∞.

#### Lemma 2.

*A necessary and sufficient condition for S to lie in the consistency zone for R*\* *with sequence distances under the MSCR-JC(k) model is that for all 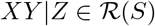*,

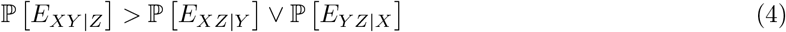

*Here* 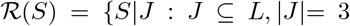, and *S*|*J is binary*} *is the set of restricted rooted triples of S (see [SS^+^03]).*

See Appendix A for a proof.

By Lemma 1, with probability one, an ancestral recombination graph 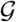 generated according to the MSCR process has the property that sequences of increasing length *k* generated on it by the Jukes-Cantor process satisfy the almost sure limit 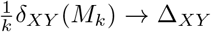 as *k* → ∞. Since almost sure convergence implies convergence in distribution, it holds that under the joint process which combines both genealogical and mutational processes, 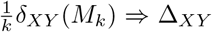 as *k* → ∞ for all *X, Y* ∈ *L*(*S*). Therefore, since the distribution function of Δ_*XY*_ is continuous, ℙ[*E*_*XY* |*Z*_] → ℙ [*E*] and ℙ[*E*_*Y Z*|*X*_] → ℙ [*F*] as *k* → ∞, where *E* := [Δ_*AB*_ < Δ_*AC*_ ∧ Δ_*BC*_] and *F* := [Δ_*BC*_ < Δ_*AB*_ ∧ Δ_*AC*_]. Therefore inequality (3) will hold for sufficiently large *k* provided that

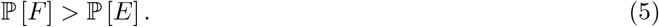

We detail the proof next.

### 2.2 Key Lemmas

Since recombination occurs only in population *A*, the number of recombination events is governed by the recombination rate *ρ* and the duration *τ*_*A*_ of population *A*. The following lemma, proved in the appendix, shows that *τ*_*A*_ can be chosen sufficiently small that with high probability, zero or one recombination occurs. In the next lemma and what follows, set intersection is denoted with product notation, i.e. *XY* = *X* ∩ *Y* for events *X, Y*, and the important events are

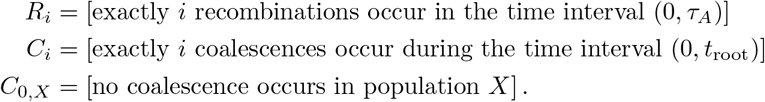

#### Lemma 3

(Recombination Probabilities). 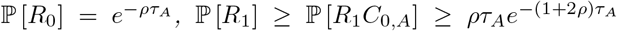, *and* 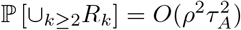 *as τ_*A*_* → 0^+^.

For the case where no recombination occurs, the probabilities of *E* and *F* are estimated in the following lemma using elementary MSC calculations which can be found in the appendix.

#### Lemma 4

(No Recombination Case). ℙ[*E*|*R*_0_] − ℙ[*F* |*R*_0_] ≤ *τ*_*AB*_.

For the case where *exactly one* recombination occurs, the following lemma characterizes the behavior of coalescent events occurring below the root of *S*. Intuitively, it says that coalescence in population *AB* is rare when *τ*_*AB*_ is small; for a detailed proof see the appendix.

#### Lemma 5

(Effect of Small Internal Edge). *As τ*_*AB*_ → 0^+^, ℙ[*C*_0_|*R*_1_] = *K* +*O*(*τ*_*AB*_), ℙ[*C*_0,*A*_|*R*_1_*C*_1_] = *O*(*τ*_*AB*_), *and* ℙ[*C*_2_|*R*_1_] = *O*(*τ*_*AB*_), *where K* = ℙ[*C*_0*,A*_|*R*_1_] ∈ (0, 1) *depends only on τ_A_ and ρ, and satisfies* 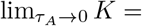 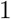 *for any fixed ρ* > 0.

Next we apply Lemma 5 to show that ℙ[*E*|*R*_1_*C*_1_] − ℙ[*F* |*R*_1_*C*_1_] is small, tending to zero as *τ*_*AB*_ → 0^+^.

#### Lemma 6.

ℙ [*E*|*R*_1_*C*_1_] − ℙ [*F* |*R*_1_*C*_1_] = *O*(*τ*_*AB*_) *as τ*_*AB*_ → 0^+^,

*Proof Sketch*. By a symmetry argument similar to that given in the proof of Lemma 4, one can show that conditional on 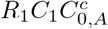, the events *E* and *F* are equally likely. Therefore by the Law of Total Probability,

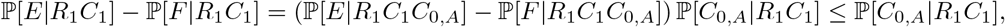

and the right-hand side is *O*(*τ*_*AB*_) by Lemma 5.

We now come to a key part of the calculation: the event *R*_1_*C*_0_, depicted in Figure 3. The next lemma demonstrates that as long as *θ* is not too large, conditional on *R*_1_*C*_0_, the event *F* is more likely than *E*.

#### Lemma 7.

*The quantity* 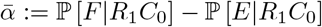 *depends only on θ and is positive if θ* ∈ (0, 3/4).

*Proof Sketch.* We sketch the proof idea here; the proof can be found in the Appendix. Conditional on *R*_1_*C*_0_, four distinct lineages enter population *ABC* at time *t*_root_. Denote these lineages by *A*_1_, *A*_2_, *B*, and *C*, as shown in Figure 3. Since no recombination occurs in population *ABC*, the order in which the lineages coalesce determines a *labeled history* (an ultrametric rooted binary tree with labeled tips and internal nodes rank-ordered according to age [RELY20]), whose tips are taken to be the lineages *A*_1_, *A*_2_, *B* and *C* at time *t*_*root*_. There are 18 such labeled histories *γ*_1_, … , *γ*_18_. Since pairs of lineages coalesce uniformly at random under the coalescent, 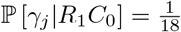 for all *j*, and hence

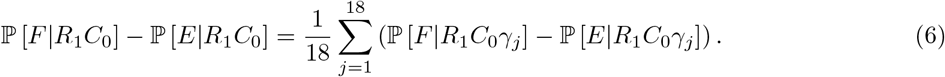

Having conditioned a particular labeled history, the probabilities 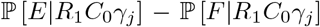 for *j* = 1, … , 18 are computed in a straightforward manner, so that the right hand side of (6) is positive provided that not too much signal is lost by a high mutation rate. In particular, since there are *two* lineages from *A* and only one from each of *B* and *C*, at least one of the *A* lineages is more likely to be included in the final coalescing pair, favoring greater pairwise distances between *A* and the other two taxa than those between *B* and *C*.

The next lemma applies Lemmas 5, 6, and 7 to show that *P* [*F*|*R*_1_] > ℙ [*E*|*R*_1_] when the internal branch length *τ*_*AB*_ is small and the mutation rate *θ* is not too large.

#### Lemma 8.

*If θ* ∈ (0, 3/4), *then* 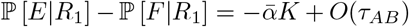 *as τ*_*AB*_ → 0^+^ *(where the term* −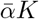 *does not depend on τ*_*AB*_).

*Proof.* By Law of Total Probability, since *R*_1_*C*_0_, *R*_1_*C*_1_, and *R*_1_*C*_2_ partition *R*_1_,

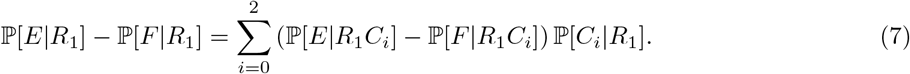

By Lemma 7, 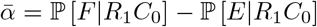 is positive and does not depend on *τ*_*AB*_. Moreover, by Lemma 5, *K* = ℙ[*C*_0*,A*_|*R*_1_] does not depend on *τ*_*AB*_ and ℙ[*C*_0_|*R*_1_] = *K* + *O*(*τ*_*AB*_), hence

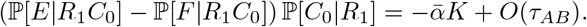

It remains to show that the other two summands on the right hand side of (7) are of magnitude at most *O*(*τ*_*AB*_). Indeed, the *i* = 1 term is at most *O*(*τ*_*AB*_) by Lemma 6. Moreover, by Lemma 5, ℙ [*C*_2_|*R*_1_] = *O*(*τ*_*AB*_), so the *i* = 2 term is at most *O*(*τ*_*AB*_) as well.

### 2.3 Proof of Theorem 1

*Proof of Theorem 1.* It suffices to prove (5) for some choice of parameters *ρ, θ, τ*_*A*_, and *τ*_*AB*_. Let *ρ* > 0 and *θ* ∈ (0, 3/4) be arbitrary; we will show that *τ*_*A*_, and *τ*_*AB*_ can be chosen sufficiently small that (5) holds. Conditioning on the number of recombination events in population *A*,

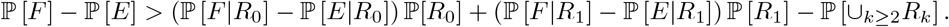

Therefore by Lemma 4 and the trivial inequality ℙ [*R*_0_] ≤ 1,

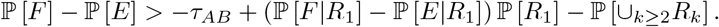

By Lemma 8, there exists *δ* > 0 such that 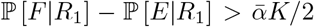 whenever 0 < *τ*_*AB*_ < *δ*. Assume further that *τ*_*AB*_ ∈ (0, *δ*). Then

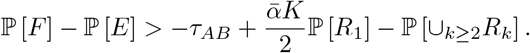

By Lemma 3, there exists constants *C, D* > 0 not depending on *τ*_*AB*_ such that 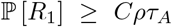 and 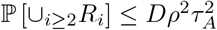, so that

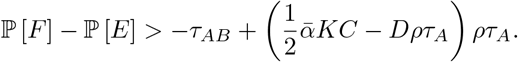

Since *K* does not depend on *τ*_*AB*_ and *K* → 1 as *τ*_*A*_ → 0 by Lemma 5, there exists *τ*_*A*_ > 0 sufficiently small that both *K* > 1/2 and 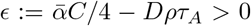. It follows that ℙ [*F*] − ℙ [*E*] > −*τ*_*AB*_ + *∊ρτ*_*A*_. Since *∊* does not depend on *τ*_*AB*_, it follows that ℙ [*F*] − ℙ [*E*] > 0 for *τ*_*AB*_ sufficiently small.

## 3 Simulation Study

We performed a simulation study to characterize the inconsistency zone established in Theorem 1. Code and documentation can be found at https://github.com/max-hill/MSCR-simulator.git. In all simulations, sequence data is generated according to the MSCR process on an ultrametric species phylogeny *S* with three species *A*, *B*, *C*, and rooted topology *AB*|*C*. In all cases, *k* = 500, *τ*_*A*_ = 1 and *θ* does not vary among populations. We use the notation 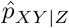 to denote the proportion of the *m* samples from which the rooted triple *XY*|*Z* was inferred, and 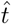 to denote the *R*\* uniquely favored rooted triple of the *m* samples. By the strong law of large numbers, 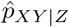 serves as an estimate of ℙ [*E*_*XY* | *Z*_] for large *m* where *E*_*XY* | *Z*_ is defined as in Lemma 2.

The range of recombination rates considered in these simulations are comparable to those in [LK12], who suggest they encompass biologically plausible values. As for mutation rates, typical rates in eukaryotes are on the order of *μ* = 10^−9^ to 10^−8^ per site per generation [Hah18, Lyn10] and effective eukaryotic population sizes *N*_*e*_ range from 10^4^ to 10^8^ [LM15], making the values considered here of *θ* = 2*N*_*e*_*μ* ∈ {0.01, 0.1} plausible as well. Computational constraints limited the ability to consider mutation rates lower than these, as doing so would have necessitated an increase in *k* or *m* to compensate; however the analytic results here predict that the inconsistency zone will persist, and may grow, for smaller values of *θ*: the computed difference 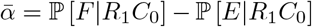 actually increases as *θ* → 0, as shown in Figure 10 in the appendix, suggesting that phylogenetic conflict may be greater under regimes with smaller mutation rates than those simulated here.

In the first experiment, we simulated the MSCR-JC(k) process under a variety of parameter regimes in order to characterize the anomaly zone and evaluate the robustness of triplet-based inference in the presence of intralocus recombination. In particular *m* = 10^4^ replicates were generated independently under each parameter regime, with the aim of estimating how frequently the correct topology was inferred. The parameters used were *θ* = 0.1, *τ*_*AB*_ ∈ {0.01, 0.02, … , 0.20}, *ρ*_*A*_ ∈ {0, … , 20}, and *ρ*_*X*_ = for all *X* ≠ *A*, so that recombination occurred only in population *A*. Figure 4 shows the value of 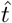 for each simulated parameter regime, and Figure 5 plots the surface 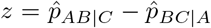 as a function of *ρ*_*A*_ and *τ*_*AB*_, so that parameter regimes with negative *z* values indicates inconsistent inference.

**Figure 4:**
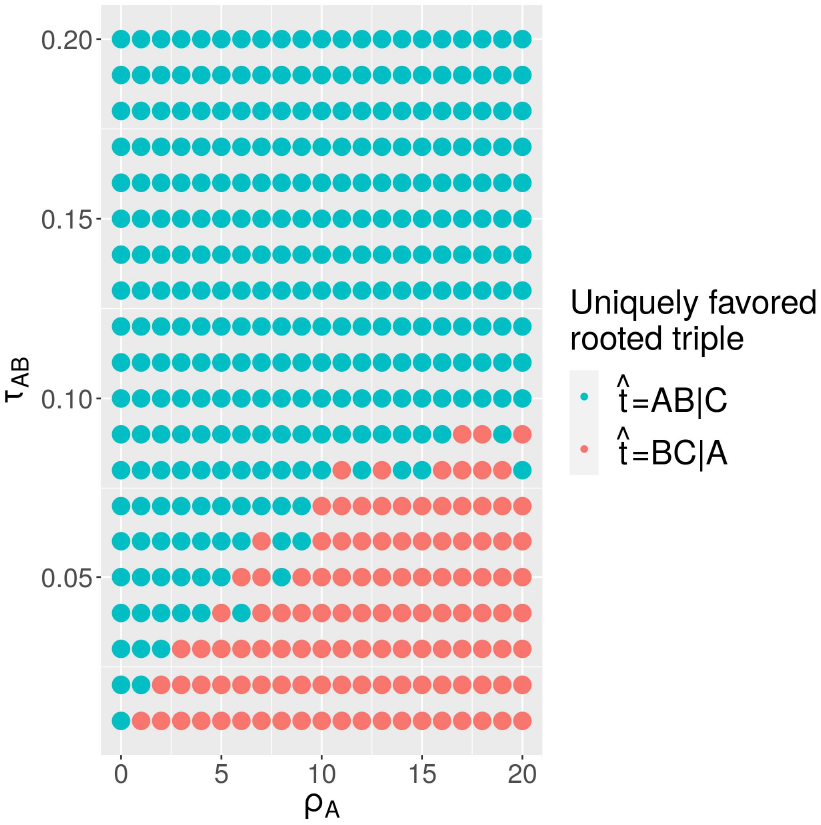
*R*\* inconsistency zone. The color of each dot represents a simulation of *m* = 10000 replicates.

**Figure 5:**
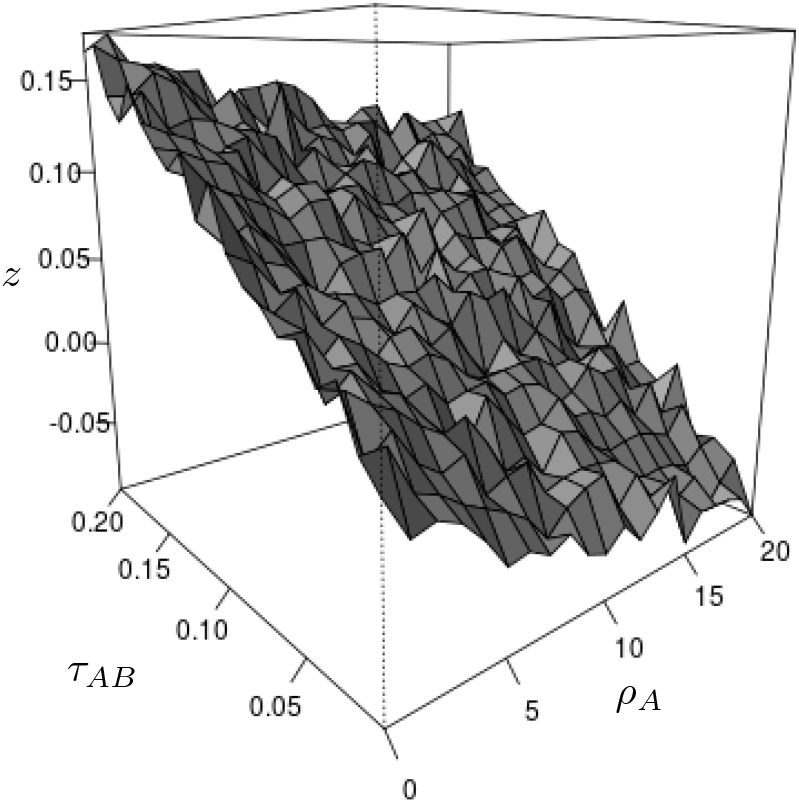
The surface 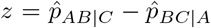 as a function of *τ*_*AB*_ and *ρ*_*A*_.

We also evaluated *R*\* inference with rooted triples inferred not by equation (1), but rather by maximum-likelihood under the (false) assumption of no intralocus recombination; in this mode, which we call ***R***\* **with maximum likelihood**, binary sequences were simulated and the maximum likelihood rooted triple was computed analytically using the method in [Yan00]. A plot almost identical to Figure 4 was obtained. For the very short internal branch length *τ*_*AB*_ = 0.01, simulations were run with similar parameters and higher number of replicates (*m* = 15, 000), with inference performed using both *R*\* with sequence distances and *R*\* with maximum likelihood. Figure 6 plots the difference 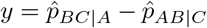 as a function of *ρ*_*A*_ obtained from these simulations.

**Figure 6:**
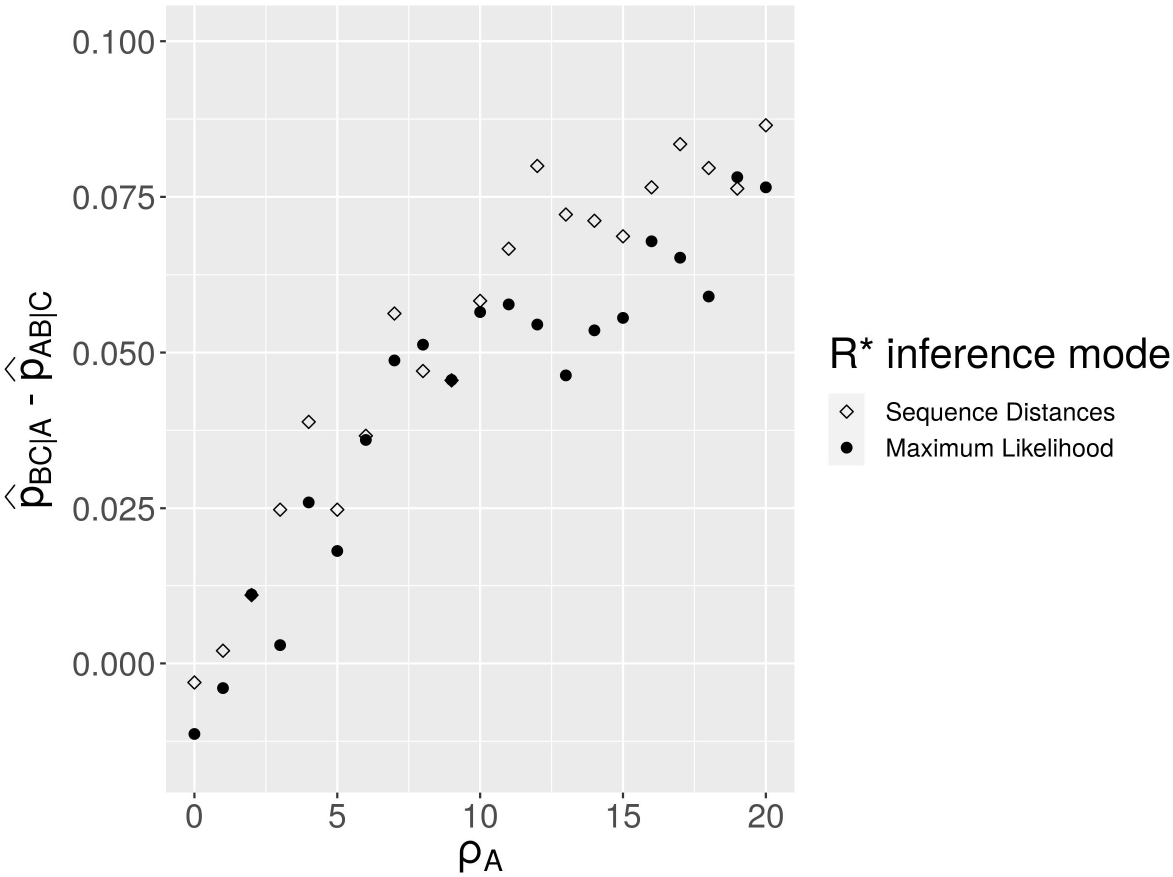
The effect of increasing *ρ*_*A*_ on inference using *R*\* with sequence distances and maximum likelihood.

These results show that the combination of intralocus recombination in population *A* along with a very short internal branch length *τ*_*AB*_ resulted in the rooted triple *BC*|*A* being more slightly likely to be inferred than the correct topology *AB*|*C*. Figure 6 shows clearly that this effect increases for larger values of *ρ*_*A*_. Nonetheless, as both Figures 5 and 6 show, the magnitude of this effect is relatively small: even when 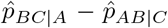 is positive, it is never greater than 0.1. Moreover, as Figures 4 and 5 show, this effect disappears when *τ*_*AB*_ is increased (ILS being less likely to occur on longer edges of *S*). Notably, even for high rates of recombination, *R*\* under both sequence distance mode and maximum likelihood mode always correctly inferred the topology of *S* when *τ*_*AB*_ > 0.1 coalescent units.

In our second experiment, we relaxed the assumption that recombination occurs only in population *A* by allowing for recombination in population *B* as well. For this simulation, *τ*_*AB*_ = 0.01 and *θ* = 0.01, with inference performed using *R*\* with sequence distances. Figure 7 shows the uniquely favored rooted triple for each choice of *ρ*_*A*_ and *ρ_B_*, with each estimate obtained from *m* = 10^5^ samples. When this experiment was repeated with *τ*_*AB*_ = 0.1, all but one parameter regimes resulted in correct inference; the exception was when *ρ*_*A*_ = 0 and *ρ*_*B*_ = 20, in which case 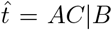. These results support the hypothesis that taxa exhibiting higher rates of recombination relative to other taxa are more likely to be inferred as more distantly related, but that the effect is small and manifests only in species triplets with very short internal branches.

**Figure 7:**
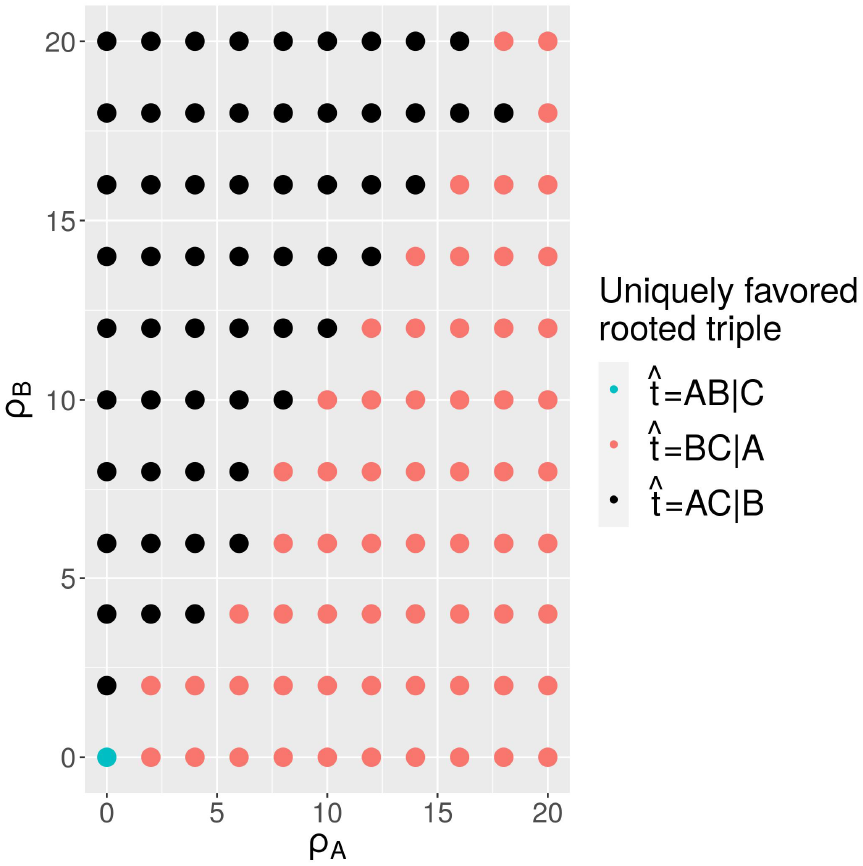
*R*\* inference with recombination in both populations *A* and *B*.

The third experiment tested the effect when all populations in *S* (excluding the root population *ABC*) experience recombination at comparable rates. The simulation parameters were *ρ* := *ρ*_*A*_ = *ρ*_*B*_ = *ρ*_*C*_ = *ρ*_*AB*_ ∈ {0, 1, … , 20} and *ρ*_*ABC*_ = 0, along with *θ* = 0.1, *τ*_*AB*_ = 0.01, and *m* = 10^6^, with inference performed using *R*\* with sequence distances. The results, shown in Figure 8, suggest that when recombination rates are similar on the edges of *S*, greater recombination rates does *not* lead to incorrect inference of rooted triples: in all cases, 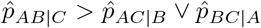, suggesting consistent inference despite the very short internal branch length, a result which agrees with the conclusions of [LK12] that even high recombination rates are not a significant source of error, at least when rates are comparable across species. Thus, the existence of differential rates of recombination between closely related taxa appears to be a necessary condition for a species tree *S* to lie in the inconsistency zone.

**Figure 8:**
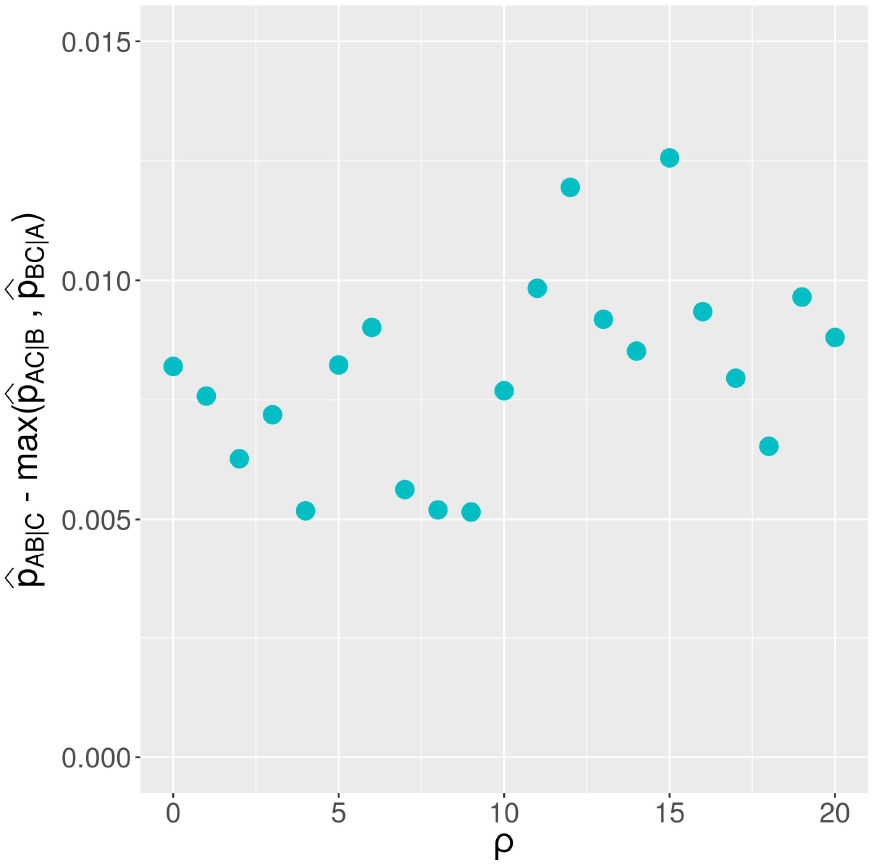
Equal recombination rates in *A, B, C* and *AB*.

## 4 Discussion

The primary focus of this study is the effect of intralocus recombination on the inference of rooted triples. In contrast to previous simulation studies [LK12, WL16, Con20], the current work considers the effect of intralocus recombination on inference of species phylogenies *with recombination rate heterogeneity across taxa*. Our main result is a proof that within the parameter space of species phylogenies there exists a subset— the inconsistency zone—in which phylogenetic conflict between the topology of the species phylogeny and the topology of inferred gene trees is of a sufficient level to render certain majority vote methods statistically inconsistent. We further quantify and characterize this inconsistency zone through simulations, showing that it includes biologically plausible recombination and mutation rates for eukaryotes, and suggesting that it arises on species phylogenies exhibiting both (1) very short internal branch lengths (less than 0.1 coalescent units) and (2) differential rates of recombination between closely related taxa. These results highlight a way in which intralocus recombination can exacerbate ILS and lead to overestimation of the divergence times of those taxa exhibiting disproportionately high intralocus recombination rates relative to other taxa.

These findings do not necessarily contradict the conclusions of [LK12] that the effect of unrecognized intralocus recombination can be minor. Indeed, our simulation experiments provide further evidence that inference of rooted triples is hampered by unrecognized intralocus recombination only in cases where the internal branch length of the species tree is short, that is in cases where ILS is already high. The size of the observed effect is also relatively small; even when the uniquely favored rooted triple do not agree with the species tree, it is usually only slightly more common than the true rooted triple. Furthermore, if differential rates of recombination between closely-related taxa are rare, then summary coalescent-based methods which take no account of intralocus recombination may nonetheless indeed be robust even when recombination rates are high.

Our results raise a number of questions for future study. Our analysis focused on a simple idealized case consisting of a rooted ultrametric three-taxa species phylogeny with mutations modeled by the Jukes-Cantor process. The nature and significance of the inconsistency zone may be affected by factors such as variable population sizes as well as elements of mutation and recombination rate heterogeneity not considered here. In addition, our theoretical results only consider distance-based gene tree estimation. Extending these results to likelihood-based inference would be of interest.

## Acknowledgments

MH was supported by supported by NSF grants DMS-1902892 (to SR) and DMS-2023239 (TRIPODS Phase II). SR was supported by NSF grants DMS-1902892 and DMS-2023239 (TRIPODS Phase II). MH and SR are grateful for the feedback from Cecile Ane and her lab members as well as Claudia Solis-Lemus.

## A Preliminary lemmas

This section contains the proof of some preliminary lemmas.

### A.1 Proof of Lemma 1

We begin with some auxiliary lemmas.

#### Lemma 9.

*Suppose* 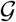 *is generated according to the MSCR process on S and* 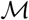 *is its associated collection of marginal gene trees. With probability one, for all* 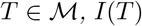, *I*(*T*) *is a finite union of nondegenerate subintervals of* [0, 1] *such that the endpoints of each of these intervals are elements of* 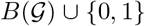 *where* 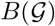 *is the set of recombination breakpoints of* 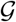.

*Proof.* Let 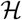 denote the collection consisting of the empty set together with all nondegenerate subintervals of [0, 1] whose endpoints are elements of 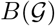, and define

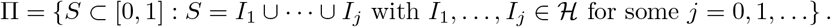

For each 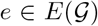 define 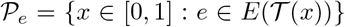. As noted in [GM97], with probability one 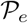 is either empty or is a finite union of intervals such that the endpoints of these intervals are a subset of 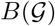. Therefore 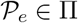 for all *e*. Let 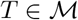 and denote the edge set of *T* by *E*(*T*). Let *x* ∈ [0, 1]. Then 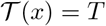 if and only if 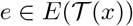 for all *e* ∈ *E*(*T*). Therefore

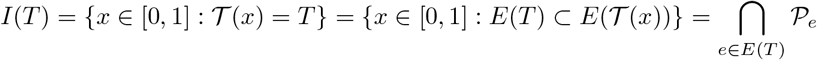

Since *E*(*T*) is finite, it will suffice to show that Π is closed under finite intersections, as the statement of the lemma will then follow from the inclusion *I*(*T*) ∈ Π. Suppose *I* = *I*_1_ ∪ … ∪ *I*_*p*_ and *J* = *J*_1_ ∪ … ∪ *J*_*q*_ with *I*_*i*_, 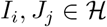 for all *i, j*. Then by the distributive property of unions and intersections,

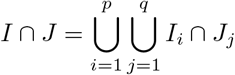

and we note that each set *I*_*i*_∩*J*_*j*_ is either empty or is an interval with endpoints in 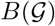. Therefore 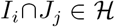 and hence *I* ∩ *J* ∈ Π. The conclusion that Π is closed under finite intersections follows by induction.

#### Lemma 10.

*With probability one, if* 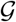 *is generated according the MSCR process on S, then*

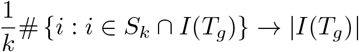

*as k* → ∞ *for all* 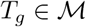.

*Proof.* We first prove the following claim: If *I* is a subinterval of [0,1] with endpoints 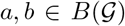, where *a* ≤ *b*, then

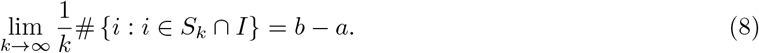

To prove this claim, first observe that

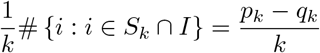

where 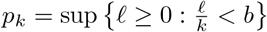 and 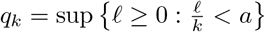 Moreover, by definition of supremum

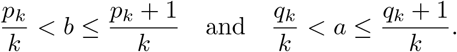

These inequalities imply that 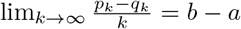. This proves the claim.

We now prove the statement of the lemma. Using the result of Lemma 9, with probability one there exists a pairwise disjoint sequence of intervals 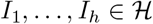 such that 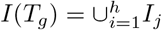. Therefore

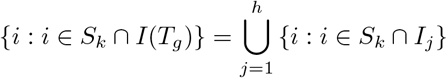

and hence

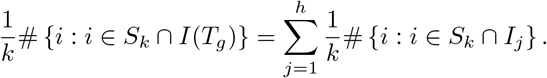

By Equation (8), 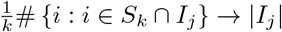 for all *j*, and hence

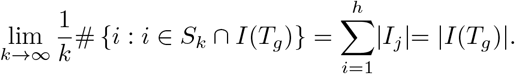

Finally, we are in a position to prove Lemma 1.

*Proof of Lemma 1.* Let *k* ≥ 2, 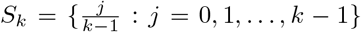, and 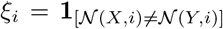, *i* ∈ *S*_*k*_. Then *ξ*_*i*_, *i* ∈ *S*_*k*_ are independent and 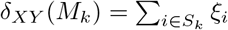. For all 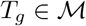,

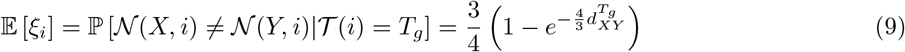

a.s. for all *i* ∈ *I*(*T*_*g*_) by definition of the Jukes-Cantor process (see e.g., [PG16]). Since 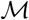 is a.s. finite,

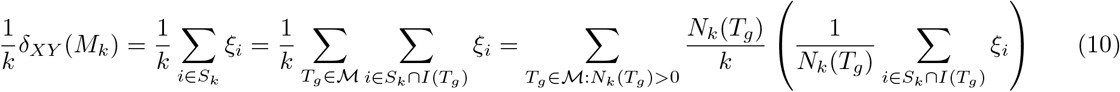

where *N*_*k*_(*T*_*g*_) = #{*i* : *i* ∈ *S*_*k*_ ∩ *I*(*T*_*g*_)}. Since *I*(*T*_*g*_) is a.s. finite union of nondegenerate intervals with endpoints in *S*_*k*_ for all 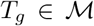, it follows that lim_*k*→∞_ *N*_*k*_(*T*_*g*_) = ∞ and lim_*k*→∞_ *N*_*k*_(*T*_*g*_)/*k* = *w*_*g*_. (By Lemmas 9 and 10.) From these limits, together with (9) and the strong law of large numbers, the right-hand side of (10) converges to Δ_*XY*_ almost surely.

### A.2 Proof of Lemma 2

*Proof of Lemma 2.* Suppose 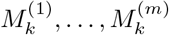 are generated independently according the MSCR-JC(k) process on *S*. For each *l* = 1, … , *m* and each triplet *X, Y, Z* ∈ *L*, let 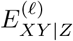 to be the event that the rooted triple inferred from *M*^(*l*)^ is *XY* |*Z*. Let 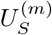 be the event that the topology of *S* is successfully reconstructed from these *m* independent samples using *R*\* with sequence distances. It suffices to show 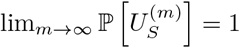 if and only if (4) holds. By definition of the *R*\* pipeline, 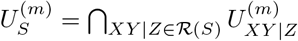 where

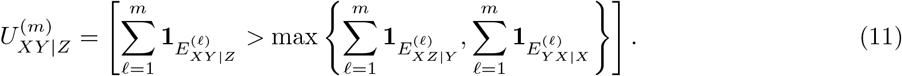

Since the samples 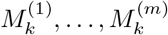 are i.i.d., the law of large numbers implies

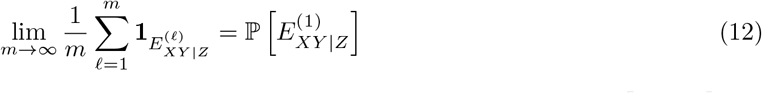

for all triplets *X, Y, Z* ∈ *L*. It follows from (11) and (12) that (4) holds if and only if 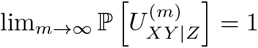 for all 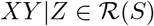. Since 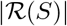 is finite, the equivalence of (4) and 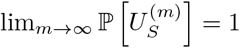 is established.

## B Key lemmas

We now prove the key lemmas leading to our main result.

### B.1 Proof of Lemma 3

*Proof of Lemma 3.* Since recombination occurs only in population *A* of *S*, the event *R*_*i*_ occurs if and only if exactly *i* recombinations occur in population *A*. Since the process starts with a single lineage in *A* at time *t*, the first event (if an event occurs in *A*) must be a recombination, which occurs with rate *ρ*. Therefore there is an exponentially distributed waiting time *e*_*ρ*_ with rate *ρ* such that a the lineage in *A* recombines at time *e*_*ρ*_ if *e*_*ρ*_ < *τ*_*A*_ and not otherwise. Therefore 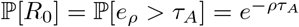.

Given *e*_*ρ*_ < *τ*_*A*_, population *A* has two lineages at time *e*_*ρ*_, so the next event (coalescence or recombination) occurs at rate 1 + 2*ρ*. Therefore there is an exponentially distributed waiting time *e*_1+2*ρ*_ with rate 1 + 2*ρ* and independent of *e*_*ρ*_ such that if *e*_1+2*ρ*_ < *τ*_*A*_ − *e*_*ρ*_ then an event occurs at time *e*_*ρ*_ + *e*_1+2*ρ*_ and not otherwise. Therefore *R*_1_*C*_0*,A*_ = [*e*_*ρ*_ < *τ*_*A*_, *e*_1+2*ρ*_ > *τ*_*A*_ − *e*_*ρ*_], and hence by the numerical inequality *e*^*x*^ − 1 ≥ *x*,

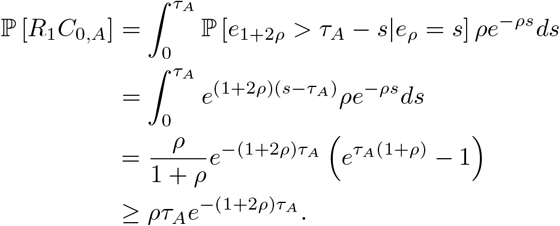

To estimate 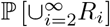, we may restrict our analysis to the single-population case since the recombination rate is positive only in population *A*. Define

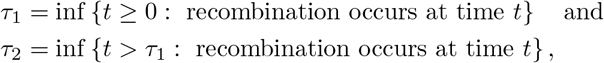

so that 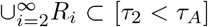. When *t* = 0, there is only one lineage in population *A*, and hence if an event occurs it must be a recombination event, which occurs at rate *ρ*. Therefore

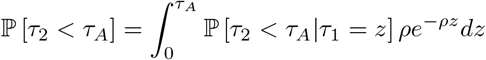

Conditional on *τ*_1_ < *τ*_*A*_, let *s*_1_ denote the time until the next event. There are two lineages at time *τ*_1_, so that events happen at rate 1 + 2*ρ*, and hence *s*_1_ ~ exp(1 + 2*ρ*), with the event at *τ*_1_ + *s*_1_ being either a recombination with probability 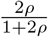 or a coalescence with probability 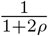. Therefore

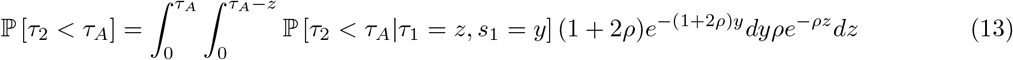

Conditional on *s*_1_ + *τ*_1_ < *τ*_*A*_, let *H*_coal_ be the event that a coalescence occurs at time *τ*_1_ + *s*_1_ rather than a recombination. Assume that *z* ∈ (0, *τ*_*A*_) and *y* ∈ (0, *τ*_*A*_ − *z*). Then by the Law of Total Probability,

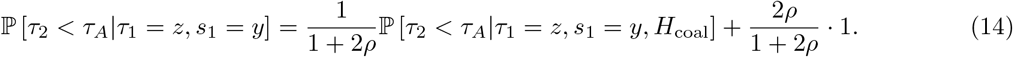

Moreover, conditional on [*τ*_1_ = *z, s*_1_ = *y*] ∩ *H*_coal_, there is exactly one lineage at time *s*_1_ + *τ*_1_ in population *A*, so the time *s*_2_ until the next event is an independent exponential with rate *ρ* and the next event must be a recombination. Therefore

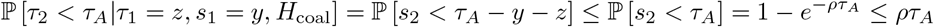

By this inequality and (14), 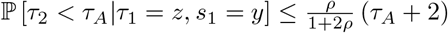. Therefore by (13),

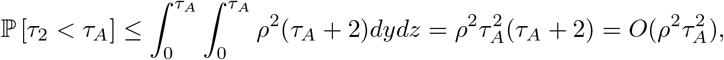

### B.2 Proof of Lemma 4

*Proof of Lemma 4.* Since the recombination rate is zero except in population *A*, it follows that conditional on *R*_0_, the ARG is a tree, so that the analysis is exactly the same as in the case of the multi-species coalescent. The lineages from *A* and *B* coalesce in population *AB* with probability 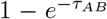. If that happens, the lineages from *A* and *B* coalesce at some time *t*_*c*_ < *t*_root_ so that

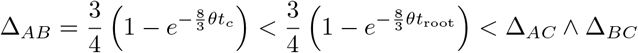

hence *E* occurs with probability one on that event. On the other hand, if the lineages *A* and *B* do *not* coalesce in population *AB*, then by a symmetry argument (each pair of lineages being equally likely to coalesce in population *ABC*), the random variables Δ_*AB*_, Δ_*AC*_, and Δ_*BC*_ are identically distributed, so that ℙ [*E*|*R*_0_*C*_0_] = ℙ [*F* |*R*_0_*C*_0_] = 1/3. It follows by the law of total probability that 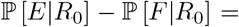 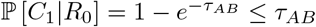.

### B.3 Proof of Lemma 5

*Proof of Lemma 5.* Since *K* ∈ (0, 1) depends only on the portion of the ARG in population *A*, it is clear that *K* does not depend on *τ*_*AB*_. By Lemma 3, 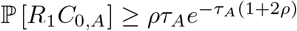 and 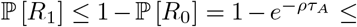 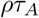. Therefore,

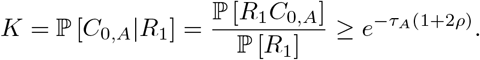

Therefore 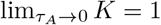 for all *ρ* > 0 uniformly in *τ*_*AB*_.

Let *e*_2_ ~ exp(1) and *e*_3_ ~ exp(3) be independent exponential clocks for the arrival of the next coalescence event given 2 and 3 lineages respectively. Then

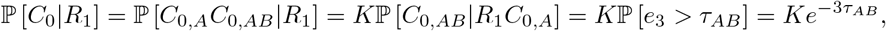

which implies

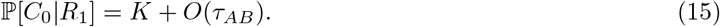

Next we claim that

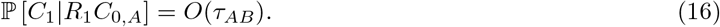

To prove this claim, observe that conditional on *R*_1_*C*_0*,A*_ three lineages enter population *AB* (i.e., two lineages from *A* and one from *B*) and one leaves (backwards in time). Thus, the event *C*_1_ occurs if and only if exactly one of the three pairs of lineages entering population *AB* coalesces and no further coalescence events occur in population *AB* (i.e., during the time interval (*τ*_*A*_, *t*_root_)). Letting 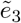 be the time until the first coalescence given three lineages, and 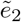 be the (independent) clock for two lineages, it follows that *C*_1_ occurs if and only if 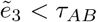 and 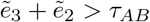. Therefore since 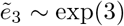 and 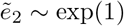,

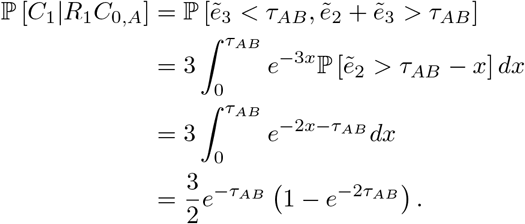

This proves (16). On the other hand, conditional on 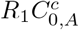, the event *C*_1_ occurs if and only if the two lineages (*A* and *B*) entering population *AB* fail to coalesce during the time interval (*τ*_*A*_, *t*_root_). Therefore

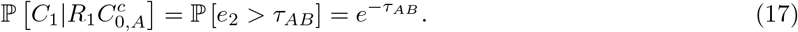

By the Law of Total Probability, 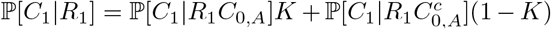, and hence by (16) and (17),

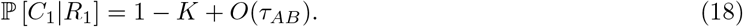

Conditional on *R*_1_, exactly one of the events *C*_0_, *C*_1_, and *C*_2_ occurs. Therefore ℙ [*C*_2_|*R*_1_] + ℙ [*C*_0_|*R*_1_] + ℙ [*C*_1_|*R*_1_] = 1. Therefore by (15) and (18), *P* [*C*_2_|*R*_1_] = *O*(*τ*_*AB*_). Finally, applying Bayes’ theorem along with the estimates from (16), (18), one obtains

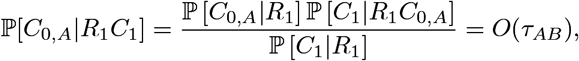

uniformly in *ρ, τ*_*A*_ as *τ*_*AB*_ → 0^+^.

### >B.4 Proof of Lemma 7

*Proof of Lemma 7.* Conditional on *R*_1_*C*_0_, four distinct lineages enter population *ABC* at time *t*_root_. We denote these lineages by *A*_1_, *A*_2_, *B*, and *C*, with *B* and *C* with the letter corresponding the originating population, as shown in Figure 3. Since no recombination occurs in population *ABC*, the order in which the lineages coalesce determines a *labeled history* (defined as an ultrametric rooted binary tree with labeled tips and internal nodes rank-ordered according to age [RELY20]), whose tips are taken to be the lineages *A*_1_, *A*_2_, *B* and *C* at time *t*_*root*_. Denote by ((*WX*)*Y*)*Z* the event that the labeled history exhibited in population *ABC* has a caterpillar topology, *W* and *X* are the first pair of lineages to coalesce, and the most recent common ancestor of *W* and *Z* coalesces with the lineage ancestral to *Y* before coalescing with that of *Z*. Denote by (*WX*)*Y Z* the event that the labeled history has balanced topology with lineages *W, X* coalescing first.

Let *t*_*k*_ be the duration of the epoch in which population *ABC* has *k* distinct lineages. Then *t*_*k*_ is an independent exponential random variable with parameter *k*(*k* − 1)/2. Conditional on *R*_1_*C*_0_, the set 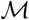 has two distinct elements, *T*_1_ and *T*_2_, where *T*_*i*_ is the triplet with leaves *A*_*i*_, *B,* and *C* for *i* = 1, 2. Hence *w*_*i*_ := |*I*(*T*_*i*_)| is uniformly distributed on [0, 1] for *i* = 1, 2 and *w*_1_ + *w*_2_ = 1. Since recombination is assumed only to occur in population *A*, it follows that 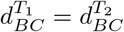, and we shall denote this quantity by *d*_*BC*_.

There are 18 labeled histories *γ*_1_, … , *γ*_18_ in which the four labeled lineages *A*_1_, *A*_2_, *B,* and *C* may coalesce. Since pairs of lineages are chosen to coalesce uniformly at random, 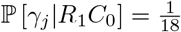 for all *j*, and hence

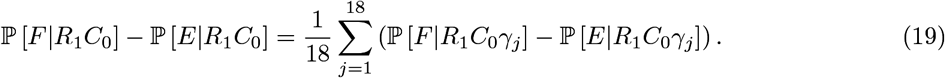

The task will therefore be to calculate ℙ [*F*|*R*_1_*C*_0_*γ*_*j*_] − ℙ [*E*|*R*_1_*C*_0_*γ*_*j*_] for *j* = 1, *… ,* 18. To do this we group together labeled histories that involve similar calculations into five claims.

#### Claim 1.

*Let γ*_1_ = ((*A*_1_*A*_2_)*B*)*C, γ*_2_ = ((*A*_1_*B*)*A*_2_)*C, and γ*_3_ = ((*A*_2_*B*)*A*_1_)*C. Then*

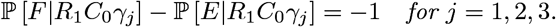

*Proof.* We prove the case for *j* = 1 as the other two cases are similar. The labeled history corresponding to *γ*_1_ is depicted in Figure 9. Since *C* is the last lineage to coalesce, 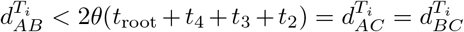 for *i* = 1, 2. Letting 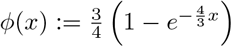, it follows that

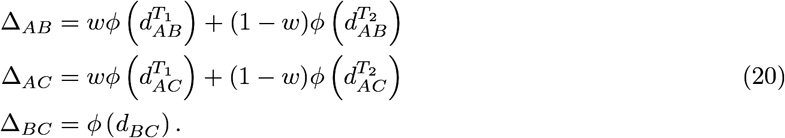

Therefore, since *φ* is increasing, Δ_*AB*_ < Δ_*AC*_ ∧Δ_*BC*_ a.s., hence ℙ [*E*|*R*_1_*C*_0_*γ*_1_] = 1. This proves the claim.

#### Claim 2.

*Let γ*_4_ = ((*A*_1_*A*_2_)*C*)*B, γ*_5_ = ((*A*_1_*C*)*A*_2_)*B, and γ*_6_ = ((*A*_2_*C*)*A*_1_)*B. Then*

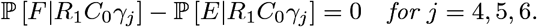

*Proof.* The proof is similar to that of Claim 1. Since *B* is the last lineage to coalesce, 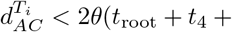 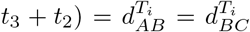 for *i* = 1, 2. Therefore Δ_*AC*_ < Δ_*AB*_ ∧ Δ_*BC*_ almost surely, so that ℙ [*E*|*R*_1_*C*_0_*γ*_*j*_] = ℙ [*F* |*R*_1_*C*_0_*γ*_*j*_] = 0. This proves the claim.

**Figure 9:**
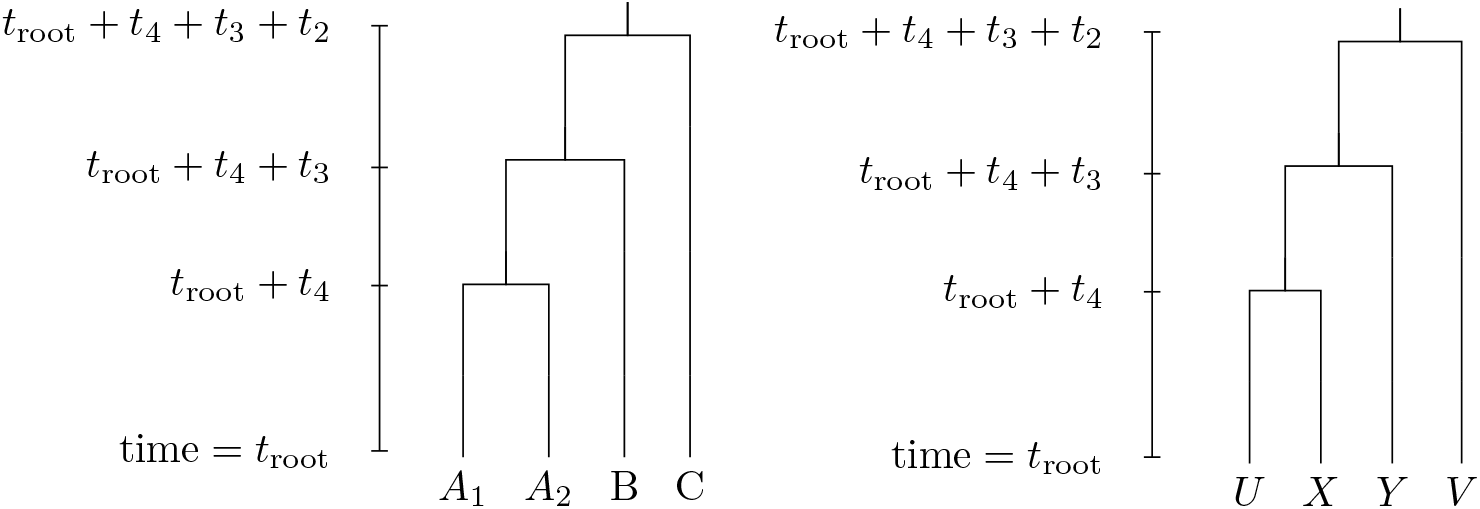
The labeled histories corresponding to Claims 1 (left) and 4 (right).

#### Claim 3.

*Let γ*_7_ = (*A*_1_*A*_2_)*BC, γ*_8_ = (*BC*)*A*_1_*A*_2_, *γ*_9_ = ((*BC*)*A*_1_)*A*_2_, *and γ*_10_ = ((*BC*)*A*_2_)*A*_1_*. Then*

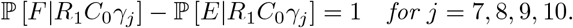

*Proof.* It suffices to show Δ_*BC*_ < Δ_*AB*_ ∧ Δ_*AC*_ a.s., conditional on *R*_1_*C*_0_*γ*_*j*_. Therefore by (20) it suffices to show that 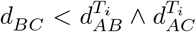 almost surely for *i* = 1, 2. Indeed, if *j* = 7, 8 then 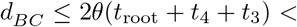 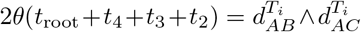, and if *j* = 9, 10 then 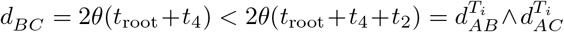. This proves the claim.

#### Claim 4.

*Let γ*_11_, *γ*_12_, *γ*_13_, *γ*_14_ *be* ((*A*_1_*C*)*B*)*A*_2_, ((*A*_2_*C*)*B*)*A*_1_, ((*A*_1_*B*)*C*)*A*_2_ *and* ((*A*_2_*B*)*C*)*A*_1_ *respectively. Then*

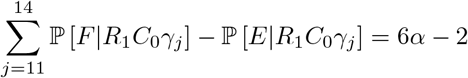

where

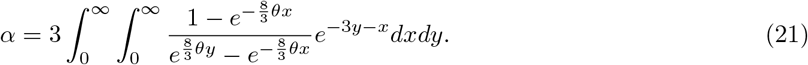

*Proof.* Consider the general case ((*UX*)*Y*)*V* where {*U, V*} = {*A*_1_, *A*_2_} and {*X, Y*} = {*B, C*}. This corresponds to the labeled history depicted in Figure 9

Let 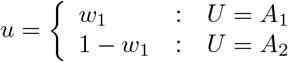, it follows that

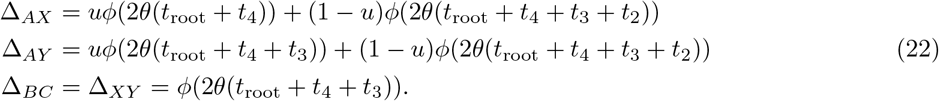

Since *φ* is increasing, Δ_*BC*_ < Δ_*AY*_ almost surely, and hence

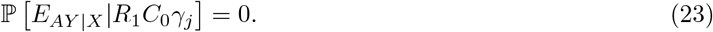

Therefore there exists some real number *α* with 0 ≤ *α* ≤ 1 such that

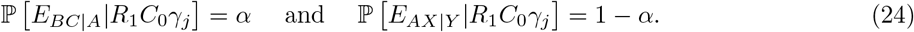

Using (22) and (24),

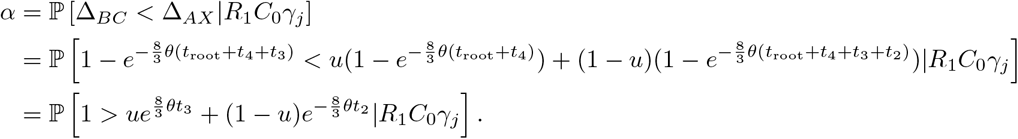

Conditional on *R*_1_*C*_0_, the random variables *t*_2_, *t*_3_ are exponentially distributed with rates 1 and 3 respectively and are independent of *γ*_*j*_ and each other. Conditional on *R*_1_*C*_0_*γ*_*j*_, *u* is uniformly distributed on (0, 1) for any *j*. Therefore

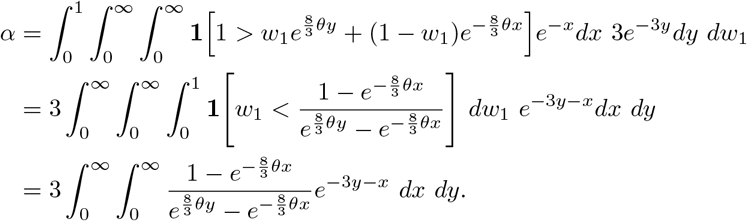

This shows that the value of *α* does not depend on *j*. Therefore by (23) and (24),

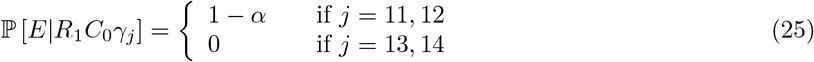

and by (24),

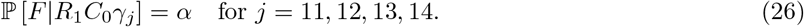

The statement of the claim then follows from (25) and (26). This proves the claim.

#### Claim 5.

*Let γ*_15_, *γ*_16_, *γ*_17_ *and γ*_18_ *be* (*A*_1_*B*)*A*_2_*C,* (*A*_2_*B*)*A*_1_*C,* (*A*_1_*C*)*A*_2_*B, and* (*A*_2_*C*)*A*_1_*B respectively. Then*

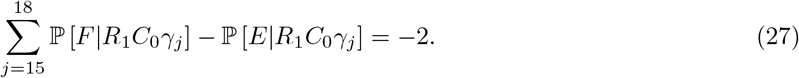

*Proof.* To prove this, we consider the general case (*UX*)*V Y*, where {*U, V*} = {*A*_1_, *A*_2_} and {*X, Y*} = {*B, C*}. This corresponds to the following labeled history:

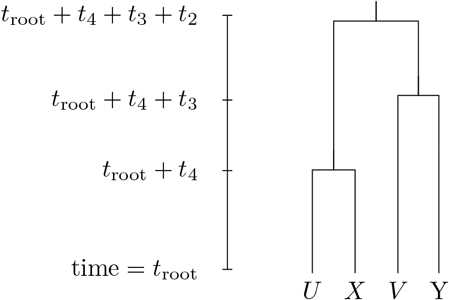

Letting 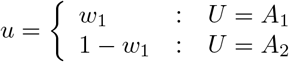, it follows that

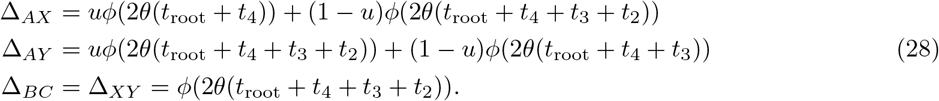

Since Δ_*BC*_ > Δ_*AX*_, it follows that

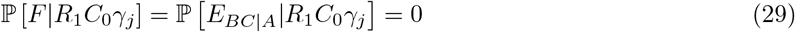

whenever 15 ≤ *j* ≤ 18. Therefore

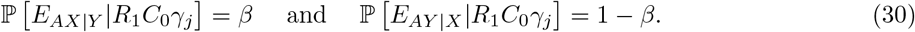

where 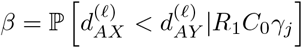. By (28),

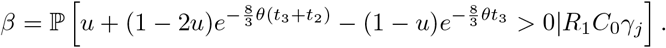

Moreover, similarly as in Claim 4, conditional on *R*_1_*C*_0_, the random variables *t*_2_, *t*_3_ are exponentially distributed with rates 1 and 3 respectively and are independent of *γ*_*j*_ and each other, and conditional on *R*_1_*C*_0_*γ*_*j*_, *u* is uniformly distributed on (0, 1) for any *j*. Therefore

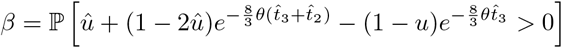

where 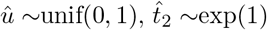 and 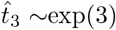, with 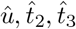 independent. In particular, this implies *β* does not depend on *j*. Moreover, equation 30) implies ℙ [*E*|*R*_1_*C*_0_*γ*_15_] = ℙ [*E*|*R*_1_*C*_0_*γ*_16_] = *β* and ℙ [*E*|*R*_1_*C*_0_*γ*_17_] = ℙ [*E*|*R*_1_*C*_0_*γ*_18_] = 1 − *β*. Therefore 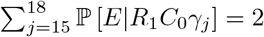. Combining this with (29) implies equation (27). This proves the claim.

We now return to the proof of the lemma. Combining the results from Claims 1-5 with (19) yields

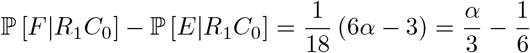

and substituting the formula for *α* from (21) gives

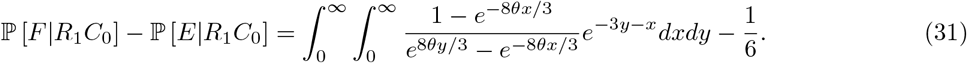

It will suffice to show that the right hand side of (31) is a decreasing function of *θ* and equals zero when *θ* = 3/4, as this will imply that it is positive for all 0 < *θ* < 3/4. By the substitution *c* = 8*θ/*3, it suffices to show that the function

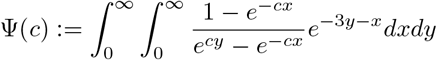

satisfies Ψ(2) = 1/6 and is strictly decreasing. Numerical plots of Ψ are given in Figure 10. Making the substitution *u* = *e*^−(*x*+*y*)^, *du* = −*e*^−(*x*+*y*)^*dx*,

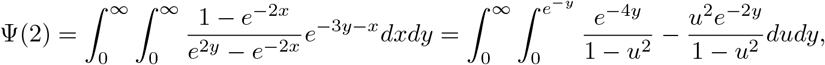

and by the Fubini-Tonelli theorem,

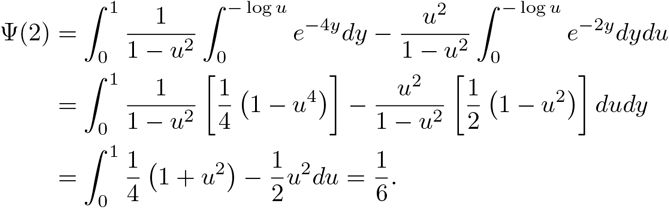

**Figure 10:**
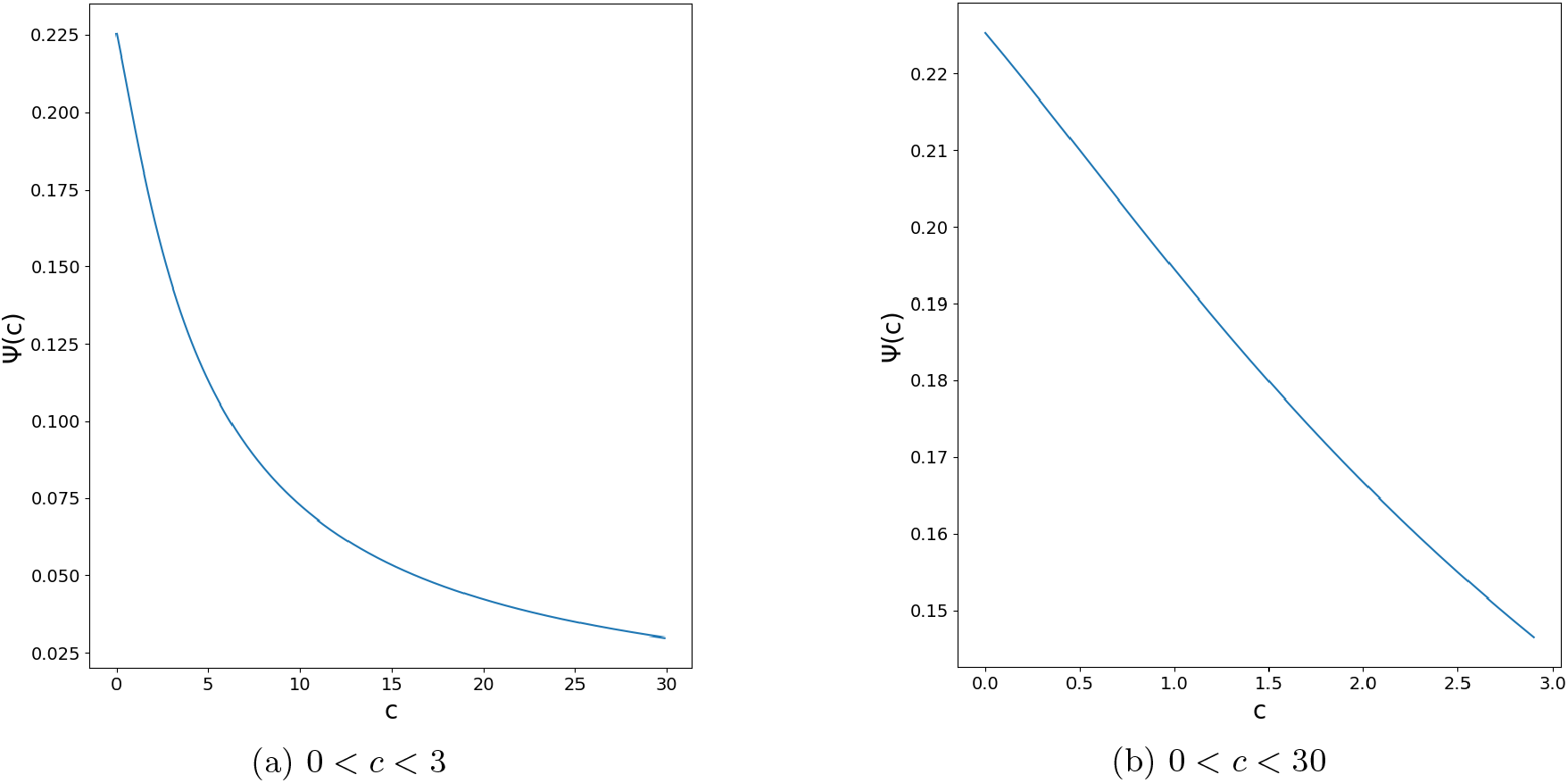
Numerical plots of the function Ψ(*c*), shown in blue. Note that Ψ is decreasing and Ψ(2) = 1/6.

It remains to show that Ψ(*c*) > Ψ(2) for all 0 < *c* < 2. Fix *x, y* > 0 and let 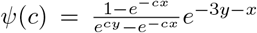. Differentiating *ψ* gives

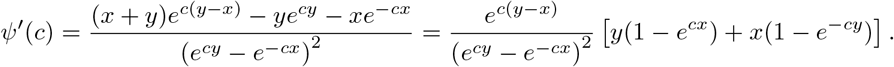

Since 1 − *e*^*a*^ < −*a* for al *a* ≠ 0, *y*(1 − *e*^*cx*^) + *x*(1 − *e*^−*cy*^) < *y*(−*cx*) + *x*(*cy*) = 0, it follows that *ψ*′ (*c*) < 0 for all *c* > 0 and hence Ψ is strictly decreasing on (0, ∞), as required.

## C Inconsistency of Unrooted Quartet Majority

Our main result extends to the unrooted case. For *W, X, Y, Z* ∈ *L*, the unrooted quartet topology inferred by the *four point method* is *WZ*|*Y Z* if *δ*_*W X*_ + *δ*_*Y Z*_ < (*δ*_*W Y*_ + *δ*_*XZ*_) ∧ (*δ*_*W Z*_ + *δ*_*XY*_). Let *S* be an ultrametric species phylogeny with *L* = {*A, B, C, D*} and topology (((*AB*)*C*)*D*). Then the unrooted quartet topology is *AB*|*CD*, but we will show that it may not be the most likely unrooted quartet topology to be inferred by the four point method when the data *M*_*k*_ is generated according to the MSCR-JC(k) process on *S* for *k* sufficiently large.

Specifically, take *S* be the tree from the proof of Theorem 1. We may append a distantly related fourth leaf *D* to an edge above the root as depicted in Figure 1b. Then the resulting tree has unrooted quartet topology *AB*|*CD*. However the residual bias in favor of the *BC*|*A* rooted triple rather than *AB*|*C* in subtree *S* causes the four point method to infer the unrooted quartet topology *BC*|*AD* more often than *AB*|*CD*. This idea motivates the following corollary.

### Corollary 1.

*For k sufficiently large, there exists a species phylogeny S with four leaves such that the most likely unrooted quartet topology to be inferred using the four-point method from MSAs generated under the MSCR-JC(k) model is different from the unrooted quartet topology of S.*

*Proof.* Let *q*_*k*_ be the unrooted quartet topology inferred from *M*_*k*_ by the four-point method. We wish to show that ℙ [*q*_*k*_ = *AB*|*CD*] < ℙ [*q*_*k*_ = *AD*|*BC*] for *k* sufficiently large. For each quartet *W, X, Y, Z* ∈ *L*, let

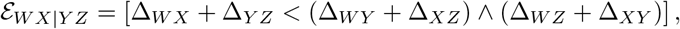

so that 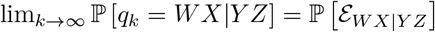 by Lemma 1. Therefore it suffices to show that 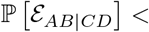 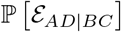.

Let *ϵ* = ℙ [*F*] − ℙ [*E*], where *E* = [Δ_*AB*_ < Δ_*AC*_ ∧ Δ_*BC*_] and *F* = [Δ_*BC*_ < Δ_*AB*_ ∧ Δ_*AC*_]. Since the restricted subtree *S*|{*A, B, C*} satisfies the assumptions of Theorem 1, it follows that for *k* sufficiently large there exists a choice of parameters *ρ*_*X*_ and *θ* = *θ*_*X*_ for all *X* ∈ {*A, B, C, AB, ABC*}, along with *τ*_*A*_, *τ*_*AB*_ such that > 0 for all *τ*_*ABC*_ sufficiently large. See Figure 1b.

Further assign *ρ*_*ABC*_ = *ρ*_*D*_ = 0 and *θ*_*ABCD*_ = *θ*. Since *S* is ultrametric, *τ*_*D*_ = *τ*_*A*_ + *τ*_*AB*_ + *τ*_*ABC*_, which is the age of the root of *S*. The only parameter of *S* which remains to be chosen is *τ*_*ABC*_. Let *z* be the age of the MRCA on the ancestral recombination graph of the lineages originating in populations *A, B,* and *C*. Since *τ*_*D*_ → ∞ as *τ*_*ABC*_ → ∞,

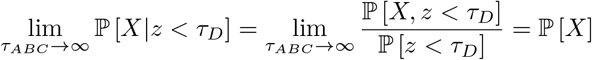

for *X* ∈ {*E, F*}. Therefore we may choose *τ*_*ABC*_ > 0 sufficiently large that

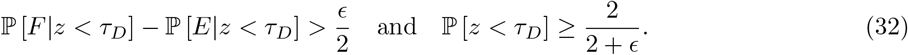

If *z* < *τ*_*D*_, then since *S* is ultrametric and *θ*_*X*_ = *θ* for all *X*, it holds that for all 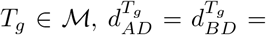 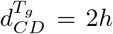 where *h* is the height of the ancestral recombination graph. Therefore Δ_*AD*_ = Δ_*BD*_ = Δ_*CD*_. Using this identity, it is easy to check that, conditional on *z* < *τ*_*D*_, the events *E* and 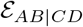 are equivalent, and the events *F* and 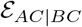 are equivalent. Therefore by the Law of Total Probability and (32),

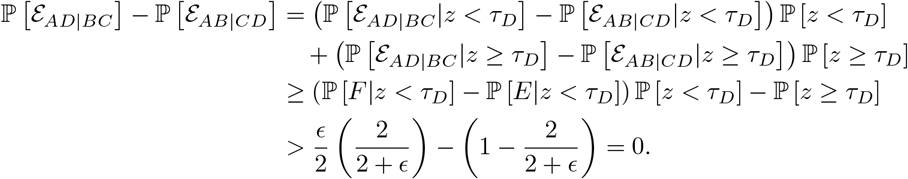

## Notes

### Competing Interest Statement

The authors have declared no competing interest.

